# A simulation-based framework to detect marine animal-vessel interactions using tracking data

**DOI:** 10.64898/2026.08.04.742798

**Authors:** David March, Leia Navarro-Herrero, Jacob González-Solís

## Abstract

Understanding how animals respond to vessels is a fundamental question in conservation, since interactions between marine animals and vessels can expose wildlife to threats such as fisheries bycatch and vessel collisions. Although animal-borne tags and vessel-tracking systems now allow both components to be monitored at fine spatiotemporal scales, distinguishing behavioural interactions from incidental co-occurrence remains challenging, particularly in areas of dense marine traffic. Here, we developed a simulation-based framework to determine whether an animal is attracted to or follows a vessel, or instead moves independently. We applied the framework to concurrent GPS data from 2,705 foraging trips by Scopoli’s shearwaters (*Calonectris diomedea*) and Automatic Identification System (AIS) trajectories from fishing and nonfishing vessels in the northwestern Mediterranean. Candidate association events (n = 1,567) were first identified using spatiotemporal proximity and subsequently tested against simulated seabird trajectories representing movement in the absence of a vessel response. The framework classified apparent associations arising when vessels approached stationary birds or when independently moving trajectories crossed by chance as independent movement. Overall, 53.5% of association events showed evidence of vessel attraction, following, or both, whereas the remainder were consistent with independent movement. Although only approximately 3% of the vessels recorded by AIS were fishing vessels, they accounted for 69% of attraction events and 74% of following events, whereas most associations with nonfishing vessels were consistent with incidental proximity. Attraction and following became less likely as birds spent more time stationary and were more likely during daytime, while following was less likely for nonfishing than fishing vessels. These ecologically coherent patterns support the biological relevance of the interactions identified by the simulation-based approach. Sensitivity analyses showed that the number of associations detected by conventional threshold methods increased markedly with broader proximity criteria, whereas inference of attraction and, particularly, following remained comparatively stable. Implemented in the R package *intersimR*, the framework provides an accessible, reproducible, and scalable approach for separating behavioural responses from incidental co-occurrence at the event level and identifying the animal-vessel interactions most relevant to conservation and management.

## 1. Introduction

In marine systems, many threats from ship-based activities, including fisheries bycatch, anthropogenic food subsidies, vessel strikes, disturbance and behavioural disruption, arise from interactions between mobile animals and vessels. Understanding how wildlife responds to vessel activity across space and time is therefore a central question in ecology, with profound implications for conservation planning and risk mitigation (Doherty et al., 2021; Ellis-Soto et al., 2023). Yet the fine-scale circumstances that lead animals to interact with vessels remain poorly understood and require further research (Hays et al., 2016; Pirotta et al., 2019; VanCompernolle et al., 2025). The increasing availability of high-resolution tracking data from both marine animals (Nathan et al., 2022; Sequeira et al., 2025; Wild et al., 2023) and vessels (Kroodsma et al., 2018; March et al., 2021; Paolo et al., 2024), offers new opportunities to quantify these interactions at unprecedented temporal and spatial detail (Jones et al., 2017; McKenna et al., 2015; Womersley et al., 2022).

Marine animals exhibit diverse behavioural responses to vessel movement, ranging from attraction to avoidance or apparent neutrality. Attraction to vessels often arises in the context of fishing activities, where discards, offal, or bait provide predictable foraging opportunities. It can also occur around nonfishing vessels when animals exploit hydrodynamic or aerodynamic advantages, such as reduced drag or uplift in the wake of the vessel (wake-riding) or the pressure wave generated at the bow (bow-riding in cetaceans). These behaviours are widespread across marine megafauna, including seabirds (Bicknell et al., 2013; Oro et al., 2013), sharks (Queiroz et al., 2019), sea turtles (Cardona et al., 2025; Casale, 2011), and marine mammals (Hamer & Goldsworthy, 2006; Luck et al., 2025). Conversely, avoidance responses have been observed in species such as cetaceans or seabirds sensitive to disturbance or ship noise (Burger et al., 2019; Erbe et al., 2019; Szesciorka et al., 2019). Importantly, not all co-occurrences reflect behavioural responses. Incidental proximity can arise when animals rest, travel independently of vessels, or move through dense traffic corridors (Clark et al., 2020; Dunlop, 2024; McKenna et al., 2015; Navarro-Herrero et al., 2025). Distinguishing attraction-driven attendance or avoidance from neutral behaviours is therefore essential for understanding potential risks, such as collision or bycatch (Lewison et al., 2004; Schoeman et al., 2020).

A variety of approaches have been developed to quantify interactions from movement data. These include methods that summarise spatiotemporal contacts between tracked individuals (Long et al., 2022), approaches that classify relative movement within dyads (i.e. paired trajectories) (Joo et al., 2021; Luisa Vissat et al., 2021), and trajectory-wide models that infer attraction or avoidance from individual movement decisions using step-selection functions (Schlägel et al., 2019). In animal-vessel studies, concurrent animal and vessel trajectories have been used to identify spatiotemporal associations across different taxonomic groups, including marine mammals (Martin et al., 2023, 2024; Mul et al., 2020; Taylor et al., 2023), but with a major focus on seabirds (Bodey et al., 2014; Cianchetti-Benedetti et al., 2018; Collet et al., 2015, 2017; Soriano-Redondo et al., 2016; Votier et al., 2010). Associations are commonly identified using predefined spatiotemporal proximity criteria, sometimes complemented by changes in animal movement or behavioural states. However, proximity-based classifications generally lack an explicit null expectation and may therefore fail to determine whether observed associations reflect directed responses. Null-model and randomisation frameworks provide a means of testing whether observed interactions depart from expected movement patterns, although their conclusions can be sensitive to how movement is represented under the null hypothesis (Chisholm et al., 2019; Miller, 2012, 2015; Périquet et al., 2021; Spiegel et al., 2016). More recently, probabilistic approaches have accounted for positional uncertainty when estimating seabird-vessel co-occurrence, focusing on the probability and duration of proximity rather than the directionality of animal movement (Rutter et al., 2024). Consequently, determining whether such associations reflect directed movement towards a vessel remains an important methodological challenge.

Among marine vertebrates, seabirds constitute a compelling model for studying animal-vessel interactions. Their foraging ecology can be strongly influenced by vessel activities, particularly fishing, which provides predictable food subsidies through discards and remains a major source of global bycatch mortality (Dias et al., 2019). At the same time, seabirds can be equipped with lightweight GPS loggers that generate high-frequency movement data, allowing detailed reconstruction of their responses to vessel trajectories (Carneiro et al., 2022; Navarro-Herrero et al., 2025; Orben et al., 2021). Accordingly, research has predominantly focused on interactions with fishing vessels using vessel tracking systems such as VMS and AIS (Clark et al., 2020; Collet et al., 2018; Collet & Weimerskirch, 2020; Corbeau et al., 2021; Orben et al., 2021; Ouled-Cheikh et al., 2020; Soriano-Redondo et al., 2016). However, seabirds also encounter nonfishing vessels, including recreational and cargo vessels, which may cause disturbance or behavioural disruption (Burger et al., 2019; Fliessbach et al., 2019; Marcella et al., 2017; Navarro-Herrero et al., 2025; Schwemmer et al., 2011; Velando & Munilla, 2011; Wong et al., 2018). Considering the full spectrum of vessel traffic is therefore important for contextualising responses to fishing vessels. However, it substantially increases the number of potential co-occurrences detected in Automatic Identification System (AIS) data, particularly in regions with dense marine traffic, making robust inference of vessel interactions especially challenging.

Here, we develop a simulation-based framework to assess dynamic animal-vessel interactions and quantify whether an animal’s movement in relation to a vessel, encompassing both attraction and potential following behaviour, deviates from random expectations. This allows us to distinguish active responses from incidental proximity (e.g. resting individuals) or chance co-occurrence with vessel trajectories (incidental crossings). Our approach assesses individual vessel association events along a seabird trajectory, providing a probabilistic estimate under two complementary null-model tests that evaluate evidence for attraction and following behaviour, respectively. The framework is implemented in an R package, enabling reproducible analyses and straightforward application to other animal-vessel tracking datasets. We illustrate its application using empirical data from concurrent GPS tracks of Scopoli’s shearwaters (*Calonectris diomedea*), a species well known to attend fishing vessels, and AIS data in a region characterised by high densities of both fishing and nonfishing vessel traffic.

## 2. Material and methods

### 2.1. Seabird tracking

#### 2.1.1. Tagging procedures

We captured breeders and nonbreeders of Scopoli’s shearwater (n = 275), using a looped pole or by hand, at their colony in Cala Morell (Menorca, Figure 1), during the breeding seasons from 2015 to 2021 (June-September). Tagging procedures followed previous studies conducted with this colony (Navarro-Herrero et al., 2024; Reyes-González et al., 2021; Soriano-Redondo et al., 2016). Individuals were fitted with different types of GPS tags (Technosmart, Italy; Perthold Engineering, U.S.A; Sextant Technology, New Zealand), that were attached at dorsal feathers using waterproof resistant Tesa tape and recorded a GPS position every 5 minutes. Total mass of devices did not exceed the recommended 4% of adult body mass (Passos et al., 2010; Phillips et al., 2003). We retrieved the tags after a minimum of 5 days after deployment and left at least one week between consecutive deployments for the same individual (number of deployments per individual and year did not exceed 3 times). In addition, we alternated tag deployments between the members of a pair to minimize impact on breeding success. Individuals were handled following relevant regulations and ASAB/ABS Guidelines for the Use of Animals in Research, with all protocols approved by the Conselleria de Medi Ambient, Agricultura i Pesca (permits: CEP24/2015, ANE06/2015, CEP30/2016, ANE08/2016, ANE24/2017, ANE27/2018, CEP21/2019, ANE19/2020, ANE18/2021).

**Figure 1.**
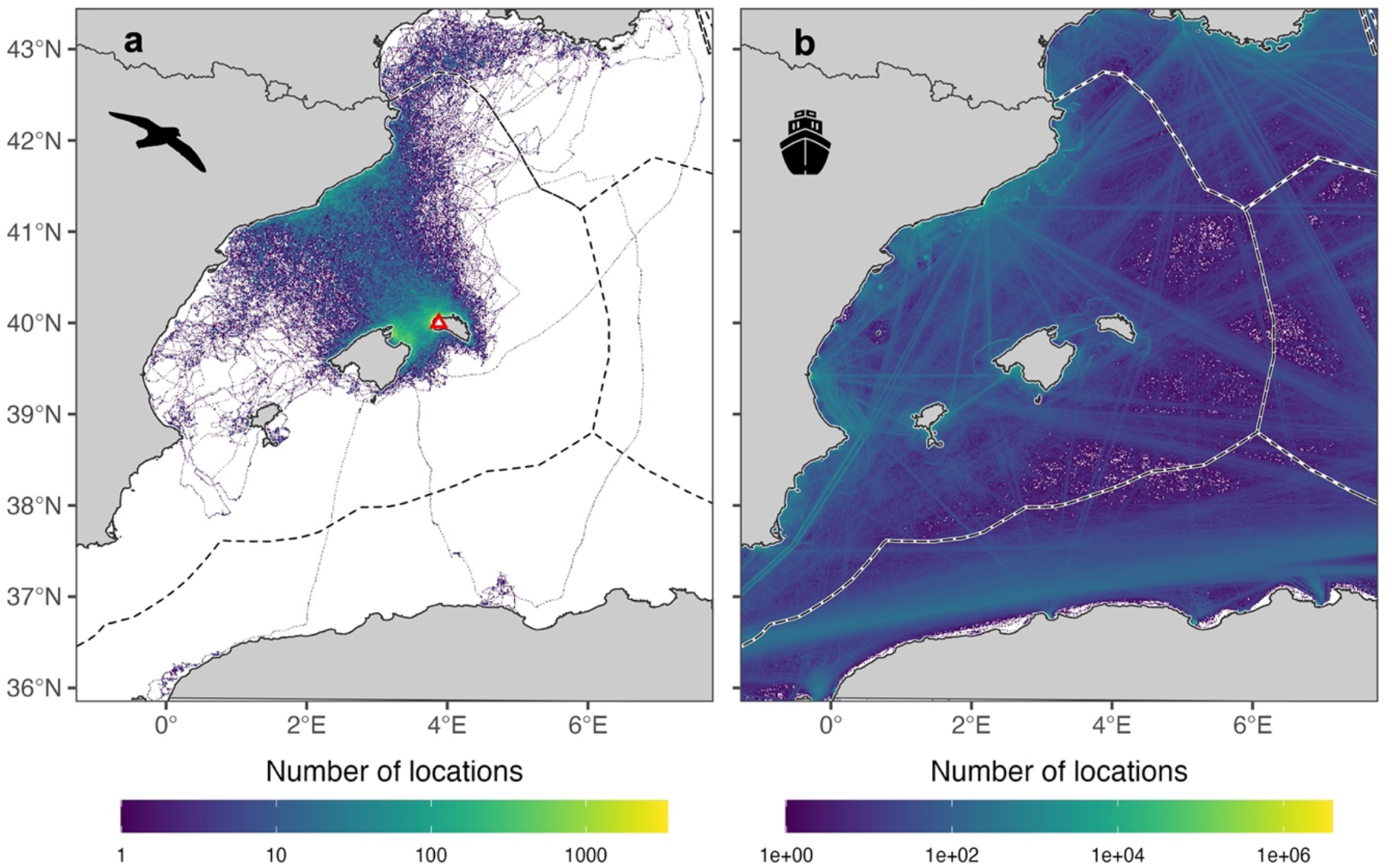
Seabird-vessel tracking data collected concurrently from 2015 to 2021 (June-September). (**a**) Density of Scopoli’s shearwater locations across all foraging trips (n = 2,705 trips). The red triangle denotes the location of their breeding colony at Cala Morell, Menorca Island. (**b**) Density of vessel locations collected from the Automatic Identification System (AIS). Dashed lines in both panels represent Exclusive Economic Zone boundaries, accessed from Flanders Marine Institute (https://doi.org/10.14284/386).

#### 2.1.2. Processing GPS data

GPS locations were filtered to remove inaccurate locations resulting from device malfunction or poor satellite reception. Near-duplicate locations, defined as animal locations that occurred 30 seconds or less after an existing location from the same animal, and which had identical longitude and latitude values were removed. GPS locations were then filtered using a speed filter that removed unrealistic movement speeds (>80 km h^-1^). We excluded short or poorly sampled trips that did not provide enough information to support inference on movement patterns. Thus, only trips with at least 20 locations, 2 hours of duration and 15 km distance were retained for further analysis. GPS tracks were then regularised at 5-min intervals using linear interpolation. Tracks with data gaps more than 2 hours were broken up for separate modelling (i.e. each portion of the track was treated independently). To characterise the movement context in which vessel interactions occurred, we applied expectation-maximisation binary clustering using the “EMbC” R package (Garriga et al., 2016; Reyes-González et al., 2021). This unsupervised approach classified each GPS location at the individual level into three different behavioural states: foraging, resting and relocating. The proportion of locations classified as resting was subsequently used as a predictor in the GLMMs assessing the drivers of vessel interactions (see section 2.3.3). Here, we refer to the “resting” state as “stationary” in subsequent analyses to emphasize its interpretation as a movement state rather than a taxon-specific behaviour.

### 2.2. Vessel tracking

Terrestrial AIS (T-AIS) data from the Western Mediterranean were collated from the Balearic Islands Coastal Observing and Forecasting System (March et al., 2021). Raw AIS data contained concurrent vessel locations to seabird tracking data at 5-min intervals (>22 million AIS messages). AIS data included individual vessel information, such as the vessel type or length. A first pre-processing of the raw data included the removal of duplicates, invalid identification numbers (i.e. Maritime Mobile Service Identity -MMSI-codes without 9 digits) and codes outside the correct numerical range (i.e. MMSI codes with first digits between 2 and 7 are those intended for individual vessels). Because we detected inconsistencies in vessel attributes associated with MMSI identifiers (e.g., changes of MMSI across years), we retained for each calendar year the more frequent combination of MMSI and vessel characteristics (e.g. vessel name and vessel type) (March et al., 2021). A common problem in AIS data is that not all fishing vessels self-report as fishing vessels (Natale et al., 2015). To identify fishing vessels from the AIS database, we matched the MMSI code against two different fishing vessel registers, the EU Fishing Fleet Register (n = 10,872 vessels with unique MMSI; https://webgate.ec.europa.eu/fleet-europa/search_en; accessed July 2024), and the Global Fishing Watch (n = 62,358 vessels with unique MMSI; https://globalfishingwatch.org/; accessed July 2024).

### 2.3. Animal-vessel interactions

To quantify seabird-vessel interactions, we followed recent works for fine-scale animal-vessel studies (Carneiro et al., 2022; Collet et al., 2017; Le Bot et al., 2018; Navarro-Herrero et al., 2025). We defined *encounter events* as instances when an animal entered a predefined detection range of a vessel (tens of kilometres), corresponding to the spatial scale at which animals may perceive and respond to vessels, and *association events* as short-distance co-occurrences (within a few kilometres) irrespective of the relative movement direction between the animal and the vessel. We further distinguished two complementary interaction processes: *attraction*, referring to directed movement towards a vessel prior to association; and *following*, referring to the persistence of an association once established. Using these definitions, we developed a two-step approach: (i) a threshold-based approach to identify encounter and association events based on the spatiotemporal co-location between seabirds and vessels, and (ii) a simulation-based framework to test whether associations were consistent with significant attraction and/or following behaviour. These processes were evaluated independently; following behaviour was assessed for all association events regardless of whether attraction was detected. The framework is implemented in the R package *intersimR* (available at https://github.com/spatialmarine/intersimR/), which provides tools for event detection and simulation-based inference, facilitating reproducible application of the method and extension to other taxa and systems.

#### 2.3.1. Threshold approach to identify seabird-vessel interactions: encounter and association events

We first identified potential seabird-vessel encounter and association events by applying spatiotemporal thresholds to paired seabird and vessel trajectories (Figure 2a). Threshold-based approaches are widely used to infer interactions from movement data (Clark et al., 2020; Orben et al., 2021) and can be further refined by incorporating behavioural information such as speed, bearing or water landings (Cianchetti-Benedetti et al., 2018; Rutter et al., 2025; Soriano-Redondo et al., 2016).

**Figure 2.**
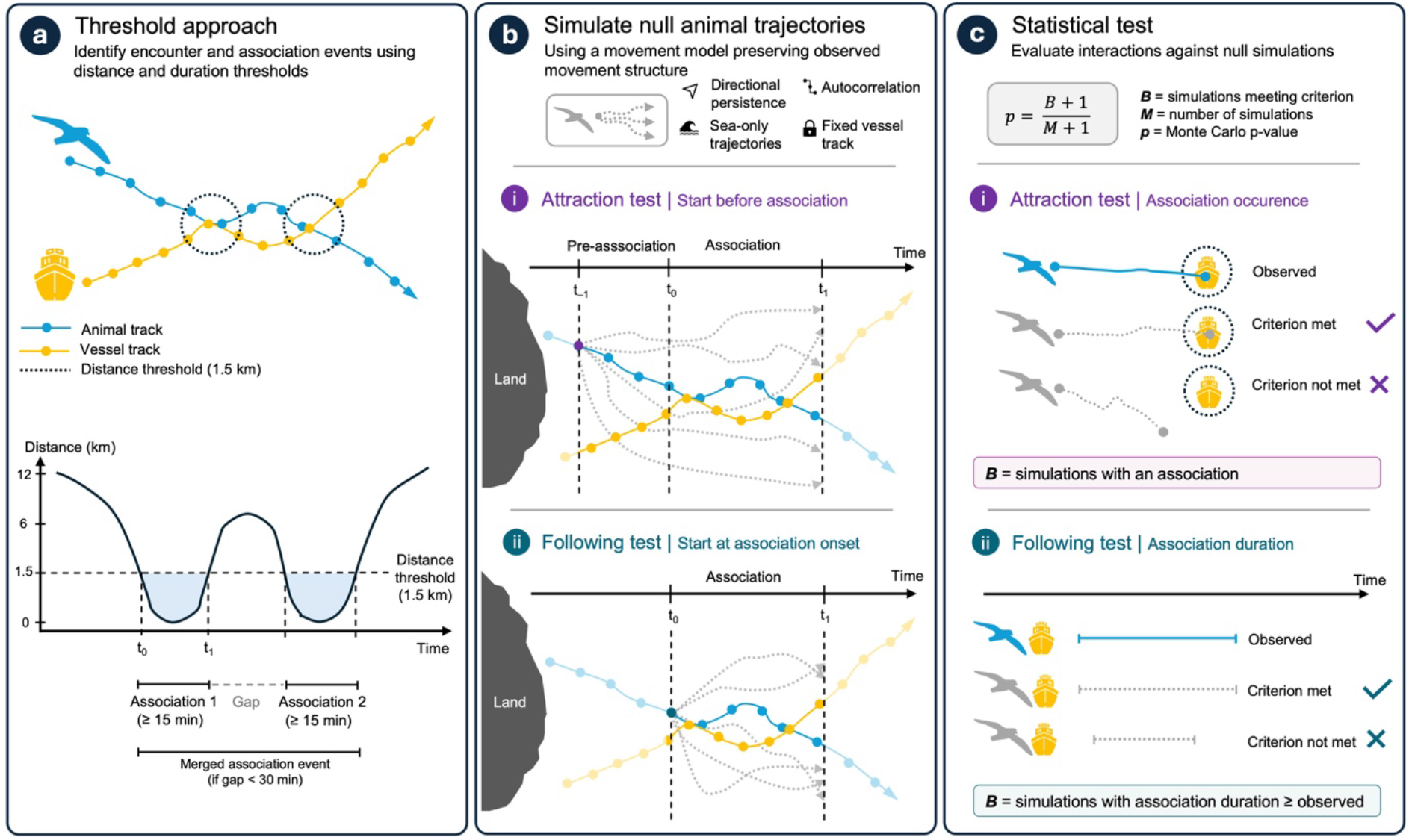
Workflow of the threshold-based and simulation-based framework used to identify animal-vessel interactions. (**a**) A threshold-based approach is first used to identify both encounter and association events from paired animal and vessel trajectories using predefined spatial and temporal thresholds. Encounter and associations occurring within a minimum temporal gap are merged into a single event. (**b**) For each association event, random animal trajectories are simulated while preserving the statistical properties of the observed movement (directional persistence and autocorrelation), constraining movements to remain at sea and keeping the vessel trajectory fixed. Simulations are initialised either before the onset of the association to evaluate attraction or at the onset of the association to evaluate following behaviour. (**c**) Observed interactions are compared against simulated trajectories using a Monte Carlo framework. For the attraction test, the test statistic is the occurrence of an association, whereas for the following test it is the duration of the association. Statistical significance is assessed as the proportion of simulations satisfying the test-specific criterion, allowing attraction and following behaviour to be evaluated independently.

Because seabird and vessel tracking systems recorded positions independently and at different sampling intervals, trajectories were first synchronised in time by linearly interpolating vessel positions to the timestamps of the seabird trajectory. No extrapolation beyond the temporal range of the vessel trajectory was performed, and interpolated positions were retained only when the nearest observed vessel fix occurred within a temporal threshold (*t_c_* = 5 min). Pairwise distances were then calculated between each seabird fix and its corresponding interpolated vessel position.

Encounter and association events were identified from the resulting seabird-vessel distance time series (Figure 2a). An event began when the distance fell below a predefined spatial threshold and ended when it exceeded that threshold. Only events exceeding a minimum duration were retained, whereas consecutive events separated by less than a predefined temporal gap were merged to avoid splitting continuous interactions caused by brief interruptions in proximity. The same event-detection algorithm was applied to identify both encounter and association events, differing only in the spatial, duration and gap thresholds used for each interaction type.

Thresholds should be selected according to both, the technical characteristics of the tracking data (e.g., location error and sampling interval) and species-specific ecological processes (e.g., sensory capabilities and movement behaviour), and may be informed by empirical observations or sensitivity analyses. To ensure that encounter and association events were temporally resolved, the minimum duration threshold was chosen to be no shorter than twice the sampling interval, thereby requiring at least two consecutive observations within the spatial threshold, following the Nyquist-Shannon sampling theorem (Nathan et al., 2022; Shannon, 1949). In this case study, encounter events were defined using a spatial threshold of 30 km, a minimum duration of 30 min, and a minimum gap of 60 min between consecutive events. The 30-km threshold corresponds to the approximate distance at which seabirds can detect fishing vessels (Collet et al., 2017). Association events were defined using a spatial threshold of 1.5 km, approximating the empirically estimated interaction distance reported for Scopoli’s shearwaters (Cianchetti-Benedetti et al., 2018). A minimum duration of 15 min (three consecutive fixes) and a minimum gap of 30 min were used to ensure robust identification of sustained association events.

#### 2.3.2. Simulation-based inference of interaction processes

Associations between animals and vessels may arise from different processes, including directed movement towards vessels (attraction) or the persistence of an interaction once established (following), as well as from random co-occurrence (e.g. incidental crossings or proximity) occurring independently of vessel presence (Collet et al., 2017; Rutter et al., 2025). To distinguish these processes from random co-occurrence, we developed a simulation-based framework to test the significance of both attraction and following behaviour. This approach evaluates the null hypothesis that seabird movements are independent of vessel presence, using simulated trajectories that preserve the movement characteristics of the observed data but are unaffected by vessel position. Although both tests rely on the same simulation procedure, they differ in their initialisation and in the metrics used to evaluate simulated outcomes. Following identification of association events (Section 2.3.1; Figure 2a), the simulation-based framework consisted of two sequential steps (Figure 2b,c):

##### Step 1. Simulating null animal trajectories

For each association event identified in Section 2.3.1, the corresponding animal trajectory was extracted from a predefined starting point until the end of the association event. Simulations were initialised using the observed position and movement characteristics of the individual at the start of this trajectory segment (Figure 2b). For the attraction test, simulations were initiated at a predefined temporal offset prior to the onset of the association, allowing evaluation of whether the movement leading to the association was directed towards the vessel. For the following test, simulations were initiated at the onset of the association, as the objective was to evaluate the persistence of the interaction once the association had already been established. The pre-association window should be sufficiently long to capture potential approach behaviour while remaining behaviourally connected to the interaction, and may vary among taxa depending on movement speeds and sensory detection ranges. For Scopoli’s shearwaters, we selected a 30-min offset, corresponding to the time required to traverse distances comparable to reported vessel detection ranges (Supplementary Methods S1). The influence of offset selection was assessed through a sensitivity analysis (section 2.3.3).

Null trajectories were generated while keeping the vessel trajectory fixed. A first-order vector autoregressive model was fitted to each observed trajectory segment using the *availability* R package (Raymond et al., 2015; Reisinger et al., 2018). This model preserves the autocorrelation and directional persistence of the observed movement. When necessary (e.g. for short trajectory segments), trajectories were linearly interpolated to provide sufficient locations (i.e. at least six) for model fitting. In addition, a minimal random displacement (jitter) was applied to avoid numerical instability in near-linear or low-mobility segments. For each association event, 1,000 null trajectories were generated, constraining movements to remain at sea using a land mask. These simulated trajectories preserve the statistical structure of the observed movement while remaining independent of vessel positions, thereby providing the null model for subsequent inference.

##### Step 2. Assessing significance of attraction and following behaviour

For each association event, we evaluated whether the observed interaction was unlikely to occur by chance under the null model described in Step 1. The simulated trajectories generated for each event were analysed using the spatiotemporal criteria defined in Section 2.3.1 (distance ≤ 1.5 km for at least 15 min), with test-specific evaluation metrics (Figure 2c). For the attraction test, we assessed whether simulated trajectories reproduced an association with the vessel. Thus, the observed statistic was the occurrence of an association, and the test evaluated whether the observed approach towards the vessel could arise under movement independent of vessel presence. For the following test, the observed statistic was the duration of the association. For each simulated trajectory, we retained the duration of the earliest simulated association; simulations in which no association occurred were assigned a duration of zero. The test then evaluated whether the observed duration exceeded that expected under the null model. A one-sided Monte Carlo p-value was computed for each test as:

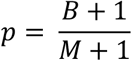

where *M* is the total number of simulated trajectories and *B* is the number of simulations in which the test-specific criterion was met. For the attraction test, *B* corresponds to the number of simulations in which an association was detected. For the following test, *B* corresponds to the number of simulations in which the duration of the earliest simulated association was equal to or greater than the observed duration. The addition of 1 in both the numerator and denominator is a standard finite-sample correction that prevents zero p-values when no simulations reproduce the observed pattern and yields an unbiased estimate of the true tail probability (Phipson & Smyth, 2010). This one-sided, non-parametric framework estimates the probability that the observed pattern would arise under movement independent of vessel presence. Events with p < 0.05 were considered significant. Significant results in the attraction test indicate directed movement towards vessels, whereas significant results in the following test indicate association persistence greater than expected under the null model.

Because the framework evaluates association events independently for each animal-vessel pair, the same period of an animal track may be associated with several vessels occurring nearby. Consequently, a movement response classified as attraction or following towards one vessel may overlap with associations involving other vessels, creating uncertainty about which vessel elicited the observed response. To assess the prevalence of this potential ambiguity, we quantified the proportion of associations that overlapped in time with associations involving other vessels for the same individual. This was calculated separately for all threshold-defined associations and for events classified as attraction, following, or both attraction and following. We also quantified the number and vessel categories of the additional vessels involved in each overlap.

#### 2.3.3. Sensitivity analyses

To evaluate the robustness of the simulation framework and its sensitivity to parameter selection, we conducted two complementary sensitivity analyses (see Supplementary Methods S2 for details). First, we evaluated how detected interaction rates varied across combinations of spatio-temporal thresholds defining association events, including proximity distance, minimum event duration, and maximum temporal gap between consecutive events. For each parameter configuration, we quantified association, attraction, and following events, and analysed interaction rates using generalised linear mixed models (GLMMs) with a negative binomial distribution, including interaction type and parameter thresholds as fixed effects, foraging trip identity as a random intercept, and trip duration as an offset term. Second, we evaluated the sensitivity of the attraction test to the temporal offset used to initialise simulated trajectories. The probability of detecting significant attraction events under different offset durations was analysed using a binomial GLMM with association identity included as a random intercept.

#### 2.3.4. Modelling predictors of attraction and following behaviours

We examined whether the probability of significant attraction and following was influenced by multiple factors, including vessel type (reclassified as fishing and nonfishing for this analysis), animal behaviour (i.e. the proportion of time the seabird remained stationary during the association event), day/night phase, and vessel density within a 30 km radius (see Supplementary Table S1 for details of each predictor). We fitted two separate generalised linear mixed models (GLMMs) with a binomial response (0 = non-significant, 1 = significant), one for the attraction test and one for the following test. In both models, individual seabird was included as a random effect. Both analyses were conducted using all association events identified by the threshold approach (n = 1,569).

## 3. Results

### 3.1. Seabird tracking

Overall, 2,705 foraging trips from 275 individuals were retained after post-processing of tracking data collected between 2015 and 2021, with annual sample sizes varying among years (Figure 1a; Supplementary Figure S2). Most of the shearwater tracking locations occurred within European waters (99.8%), mainly within Spanish jurisdiction (96.3%). The total number of trips per individual during the study period ranged between 1 and 46, with a median value of 8 trips per individual.

### 3.2. Vessel tracking

The spatial footprint of marine traffic density in the study area indicates high levels of vessel activity, particularly in coastal regions of Spain and France, as well as in the broader corridor that extends through the southern regions, connecting to the Strait of Gibraltar (Figure 1b). Most of the vessels were recreational (61%, n = 17,166) and cargo vessels (20%, n = 5,516), whereas fishing vessels accounted for only ∼3% (n = 885) of all unique vessels detected between 2015 and 2021 (n = 28,253) (Supplementary Figure S3a). The total number of monitored vessels per year remained relatively stable (8,491 ± 2,174 unique vessels per year, mean ± SD), with exception of recreational vessels that exhibited an annual growing trend (Supplementary Figure S3b).

### 3.3. Seabird-vessel interactions

#### 3.3.1. Interaction rates

The spatial distribution of encounters closely followed seabird tracking trajectories (Figures 1 and 3a). A total of 241,925 encounter events were detected across all foraging trips (Figure 3a), corresponding to an average encounter rate of 60 ± 40.9 vessels per bird per day (standardised to a 24-h period; mean ± SD, range = 1.4-308.9). The durations of encounter events, measured from the first to the last animal location assigned to each event, were strongly right-skewed, with a median of 85 min (IQR: 55-140 min). Encounters with fishing vessels accounted for 17.2% of all events (n = 41,637). In contrast, association events were more spatially constrained, occurring predominantly in coastal waters, with more than half of all events within 10 km of the coastline (median distance: 9.2 km; Figure 3b; Supplementary Figure S4). Using the threshold approach, a total of 1,567 association events were identified (Figure 3b), with an average association rate of 1.3 ± 1.1 vessels per bird per day (mean ± SD, range = 0.07-11.71). Association events were shorter than encounters overall (median = 25 min, IQR: 15-40 min), and most associations involved fishing vessels (67.2%, n = 1,054).

**Figure 3.**
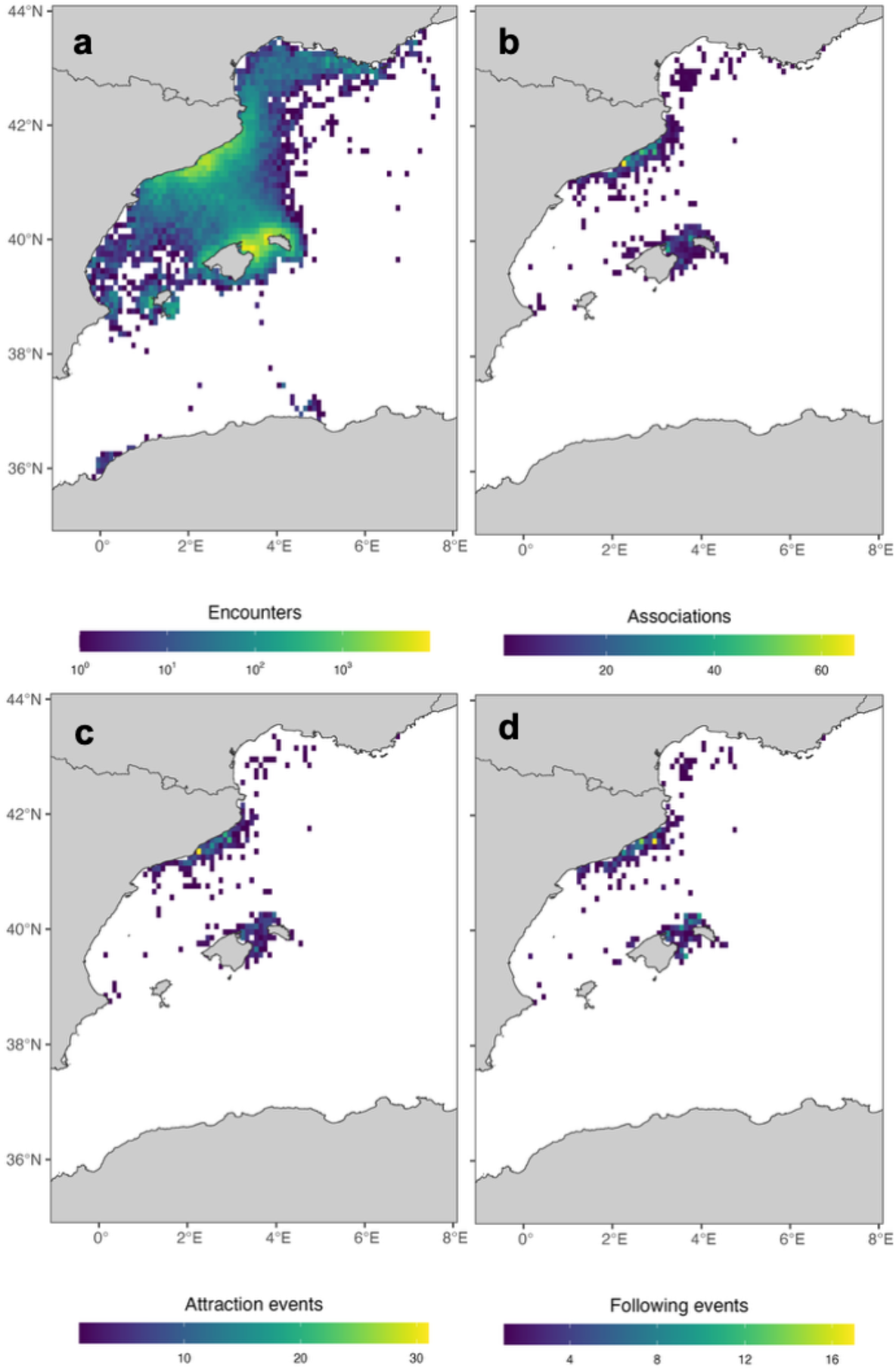
Spatial distribution and classification of seabird-vessel interaction events. (a) Density of encounters across all foraging trips (n = 241,925); note that a logarithmic (log₁₀) colour scale is used. (b) Density of associations identified using the spatiotemporal threshold approach (n = 1,567). (c) Density of significant attraction events (n = 698). (d) Density of significant following events (n = 432).

Applying the simulation-based classification, 839 associations (53.5%) were identified as significant for attraction and/or following behaviour. Attraction events accounted for 698 cases (44.5% of associations; Figure 3c). These events corresponded to an average rate of 0.89 ± 0.84 attended vessels per bird per day (mean ± SD, range = 0.07-6.86) and were predominantly associated with fishing vessels (69.1%, n = 484). Attraction events had a median duration of 25 min (IQR: 15-35 min) and were significantly longer for fishing than for nonfishing vessels (25 min vs. 20 min; W = 64,066, P < 0.001). Following events were less frequent (n = 432, 27.5% of associations; Figure 3d), with an average rate of 0.74 ± 0.65 events per bird per day (mean ± SD, range = 0.07-3.65). Similar to attraction events, following events were mainly associated with fishing vessels (74.1%, n = 320). However, they were generally longer in duration (median = 35 min, IQR: 25-60 min) and showed a stronger difference between vessel types, with longer durations for fishing compared to nonfishing vessels (45 min vs. 30 min; W = 24,174, P < 0.001).

Association events were classified into four behavioural categories according to the significance of the attraction and following tests (Figure 4). Representative examples of each category are shown together with their frequency of occurrence. Nearly half of all associations (46.5%, n = 728) were non-significant for both behaviours, indicating that most co-occurrences did not exhibit detectable responses under either test. A substantial proportion of events (26.0%, n = 407) were significant for attraction but not for following, suggesting that attraction towards vessels does not necessarily translate into sustained directional movement. Conversely, 9.0% of associations (n = 141) were significant for following but not for attraction, indicating that following behaviour can occur independently of statistically detectable attraction. Finally, 18.6% of events (n = 291) were significant for both behaviours, representing cases where individuals were both attracted to vessels and exhibited movement patterns consistent with following.

**Figure 4.**
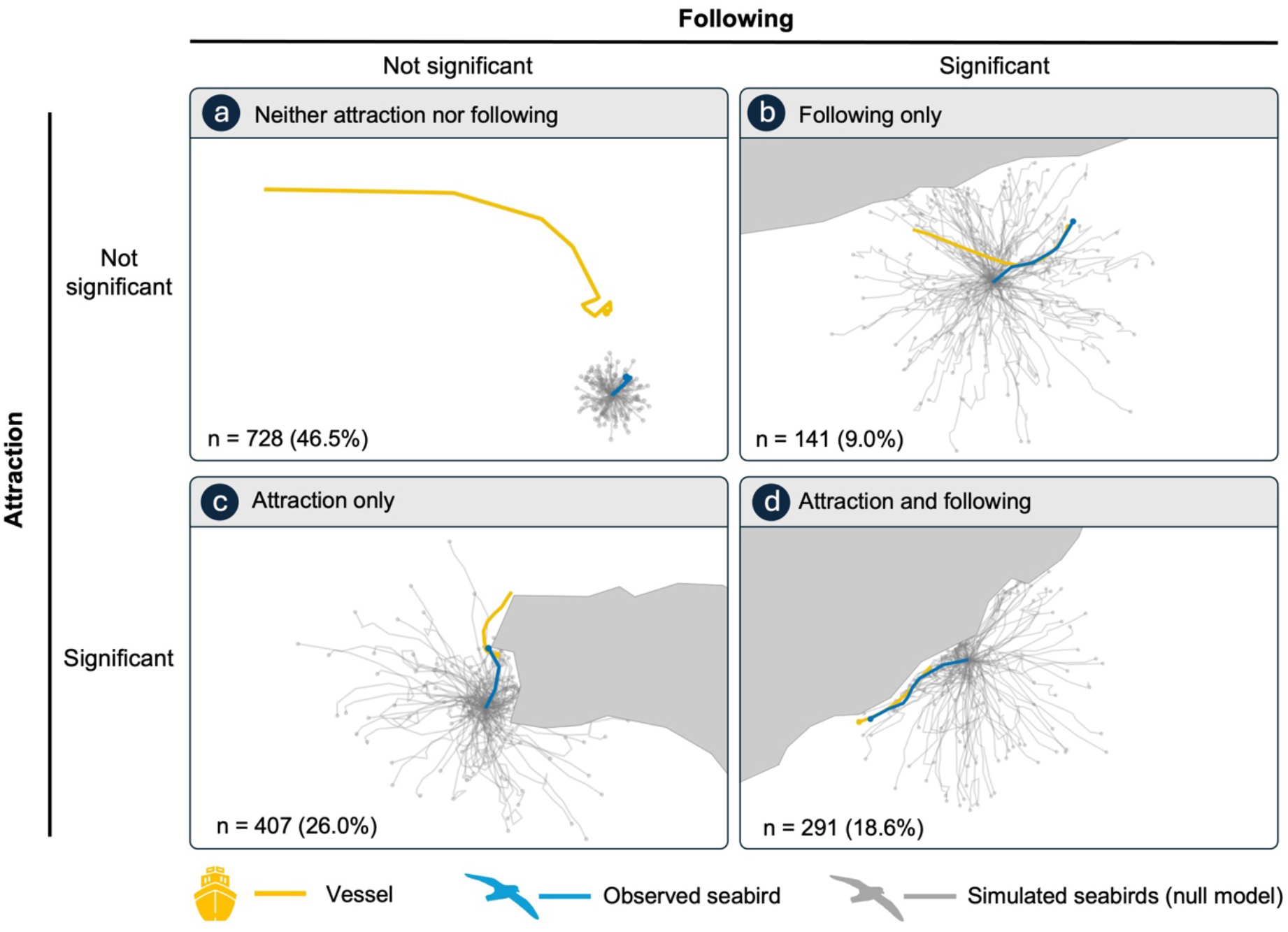
Representative examples of the four behavioural outcomes identified by the simulation framework. Panels are organised according to the significance of the attraction and following tests, illustrating interactions that were (**a**) non-significant for both behaviours, (**b**) significant for following only, (**c**) significant for attraction only, and (**d**) significant for both attraction and following. Observed seabird trajectories are shown in blue, vessel tracks in yellow, and simulated null trajectories in grey. The number and percentage of association events belonging to each behavioural category are indicated in each panel. Simulated trajectories were generated using the attraction-test initialisation to facilitate comparison of the observed approach behaviour across all four categories. Animated versions of each representative example are provided as Supplementary Movies S1-S4.

We additionally quantified associations involving multiple vessels (Supplementary Table S2). Among all associations identified using the threshold approach, 28.0% overlapped in time with an association between the same seabird and another vessel. The corresponding proportion was lower among associations classified as attraction (24.2%) or following (18.9%), and lowest among those classified as both attraction and following (13.4%). Among overlapping associations, 61.5-67.5% involved fishing vessels exclusively, depending on the interaction class considered. Within this subset, each focal association overlapped with a mean of 1.4-1.7 additional vessels, although rare cases involved up to nine.

#### 3.3.2. Sensitivity analysis

The sensitivity analysis revealed that the proximity threshold constituted the primary factor influencing interaction rates across all interaction types (Supplementary Table S3). Predicted interaction rates increased markedly with larger proximity distances, particularly for association events, for which interaction rates increased strongly at the 3-km threshold (Figure 5a). In contrast, attraction and especially following events exhibited substantially weaker responses to changes in proximity distance, as reflected by significant negative interaction terms between interaction type and proximity threshold. Minimum event duration had a moderate negative effect on interaction rates, with shorter duration thresholds generally producing higher rates across interaction types (Figure 5b). However, the magnitude of this effect was again lower for attraction and following events than for associations. By contrast, the maximum temporal gap allowed between consecutive events had minimal influence on interaction rates, with rates remaining comparatively stable across gap-time thresholds and most gap-related effects being non-significant (Figure 5c). Overall, association events exhibited considerably greater sensitivity to parameter selection than attraction and following events, which remained comparatively stable across the examined parameter space.

**Figure 5.**
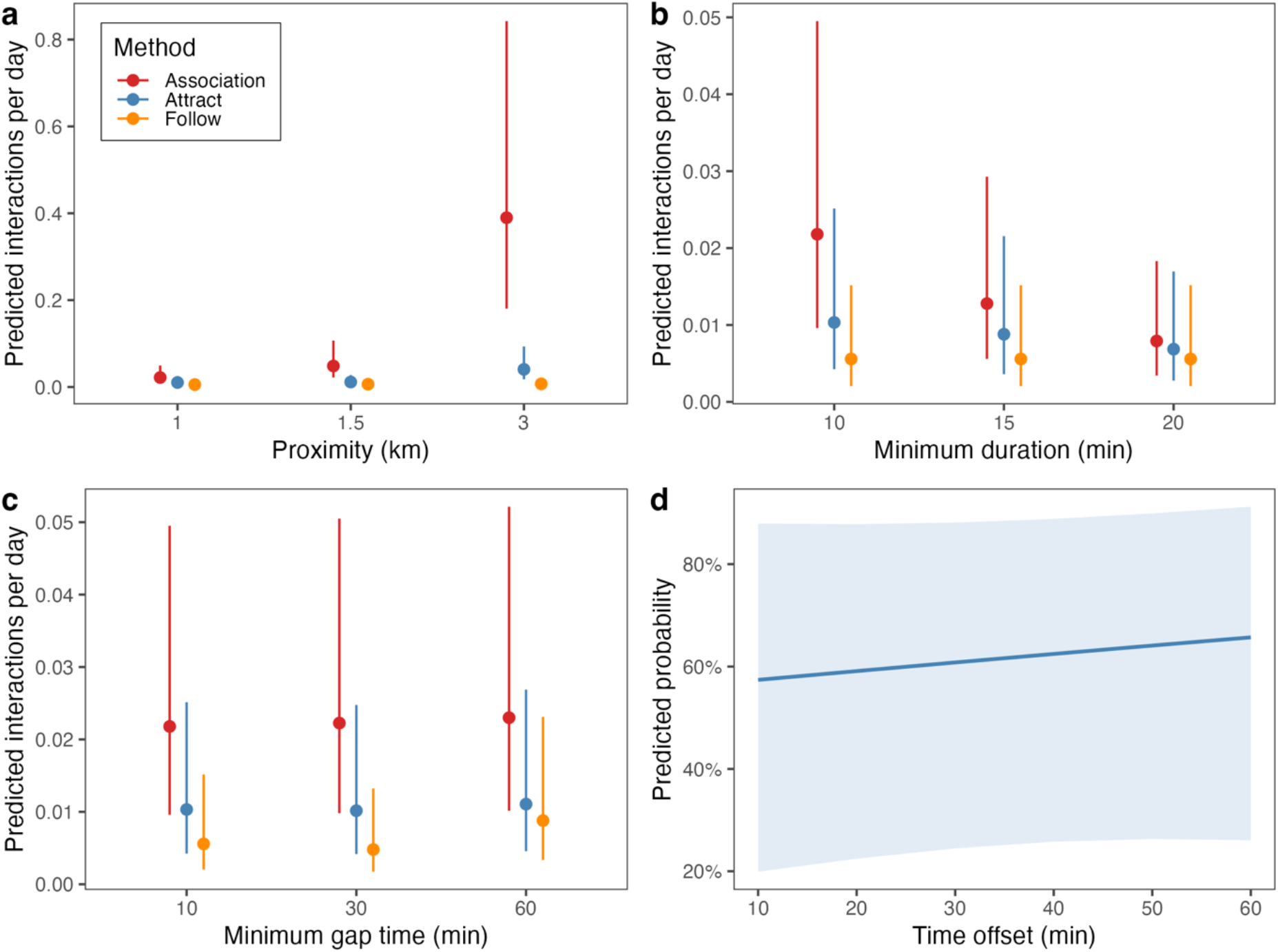
Results of the two sensitivity analyses conducted to evaluate the robustness of the interaction framework. Panels (a-c) display the partial effects of spatio-temporal thresholds on predicted interaction rates for association (red), attraction (blue), and following (orange) events. Effects shown correspond to (a) proximity distance threshold, (b) minimum event duration, and (c) maximum temporal gap allowed between consecutive events. Predicted interaction rates per day were derived from a generalised linear mixed model (GLMM) with a negative binomial distribution, including foraging trip identity (n = 50) as a random effect. Panel (d) presents the effect of varying the temporal offset used to initialise simulated trajectories in the attraction test. The plot shows the predicted probability of classifying an association as a significant attraction event across different offset durations, derived from a binomial GLMM with association identity (n = 50) included as a random effect. Shaded areas and error bars represent 95% confidence intervals.

The sensitivity analysis evaluating the temporal offset used to initialise simulated trajectories in the attraction test revealed no significant effect of offset duration on the probability of detecting significant attraction events (Supplementary Table S4). Predicted probabilities remained broadly stable across the examined range of offset values (10-60 min; Figure 5d), indicating that the attraction test was largely insensitive to moderate changes in the initialisation offset. These results support the robustness of the 30-min offset used in the main analysis.

#### 3.3.3. Factors affecting interaction significance

The generalised linear mixed models (GLMMs) revealed that the probability of an association being classified as a significant interaction varied across predictors, with both shared and contrasting patterns between attraction and following behaviours (Figure 6; Supplementary Table S5). Differences among vessel types were modest for attraction, with overlapping confidence intervals and no statistically supported effect. In contrast, vessel type influenced following behaviour, with nonfishing vessels associated with a lower probability of following compared to fishing vessels (Figure 6a). The likelihood of both attraction and following behaviours declined markedly as birds spent a higher proportion of time stationary during the association, indicating a consistent negative relationship across interaction types (Figure 6b). Time of day showed a significant effect for both attraction and following, with higher probabilities during daytime compared to night (Figure 6c). Vessel density within a 30-km radius had no statistically supported effect on either interaction type, despite a weak negative trend in both models (Figure 6d).

**Figure 6.**
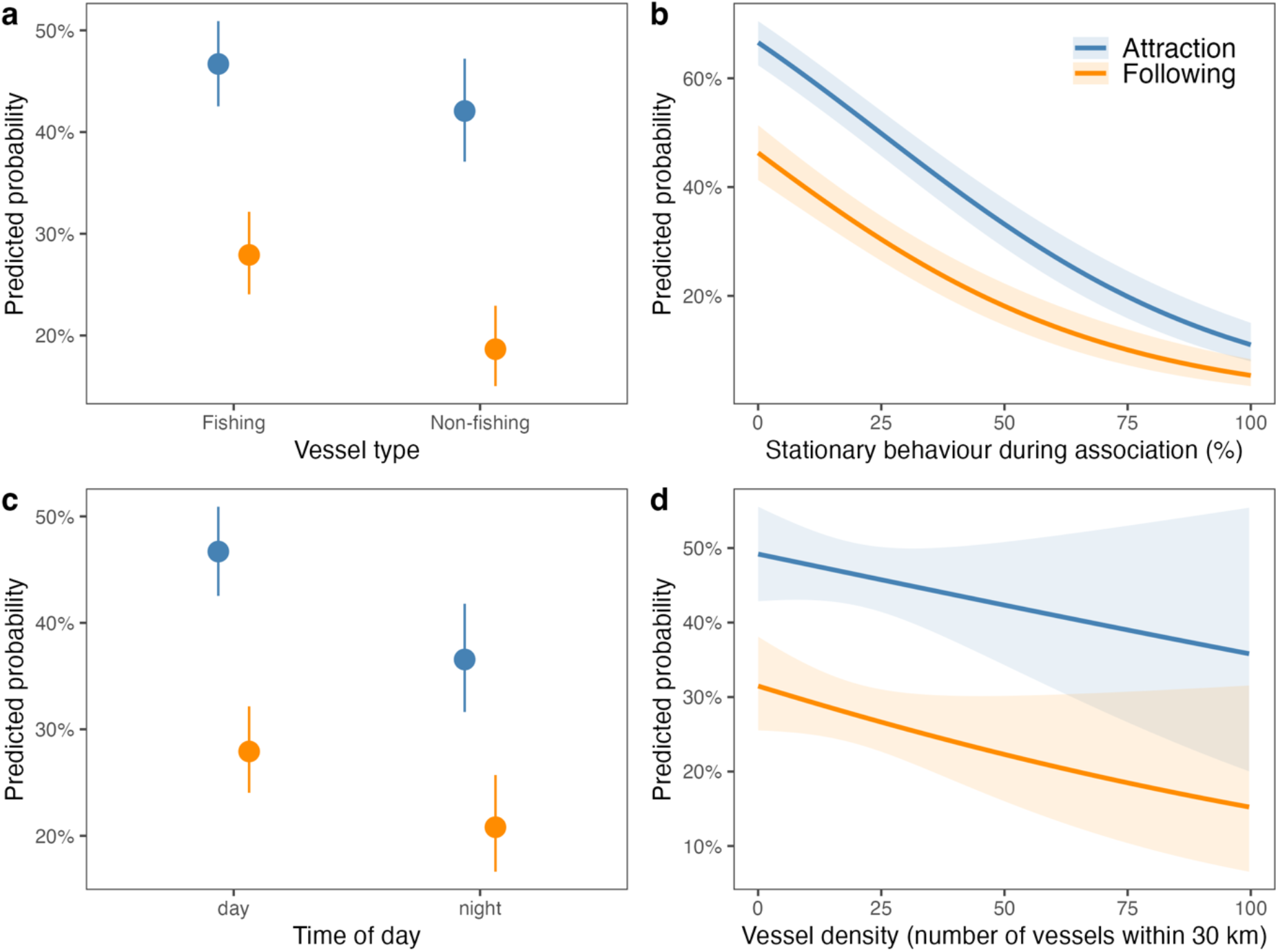
Partial effects from two independent generalised linear mixed models (GLMMs) evaluating factors associated with the probability that an association event is classified as a significant interaction, either reflecting attraction (blue) or following behaviour (orange). Shown are the modelled effects of (a) vessel type, (b) proportion of time spent stationary during the association, (c) time of day, and (d) vessel density, defined as the number of vessels detected within a 30 km radius around the association during the event. Lines and points represent fitted values, and shaded areas or error bars denote 95% confidence intervals.

## 4. Discussion

Animals and vessels may co-occur in space and time without any behavioural interaction, particularly in areas of intense marine traffic; however, active attraction and following represent a behavioural disruption usually related to anthropogenic food subsidies or energy saving and may increase the risk of bycatch or collision. Distinguishing incidental co-occurrence from directed responses is therefore critical but remains difficult using existing approaches. Here, we developed a simulation-based framework to test whether animal-vessel associations reflect attraction, following, or independent movement. Applied to Scopoli’s shearwaters, the framework classified 53.5% of associations as attraction and/or following, with fishing vessels accounting for most directed interactions despite representing only a small proportion of AIS-recorded vessels. Its relative robustness to parameter selection and the ecologically plausible patterns identified support its applicability across a wide range of animal-vessel systems.

Previous work has shown that the choice of methodological framework (i.e. whether based on co-occurrence metrics, movement classifications, or trajectory-wide models) can strongly influence inference about attraction or avoidance, particularly when behavioural states or null-model assumptions are uncertain (Chisholm et al., 2019; Miller, 2012, 2015; Périquet et al., 2021; Spiegel et al., 2016). In contrast, our simulation-based framework is designed for asymmetric human-wildlife interactions, in which animals may respond to vessels but not vice versa, and focuses on individual spatiotemporal associations. For each candidate association, the method generates null trajectories representing animal movement in the absence of a vessel response, while accounting for movement constraints and spatial barriers.

The high concentration of associations detected close to the coastline underscores the need to constrain simulated trajectories to the accessible marine domain. Preventing trajectories from crossing land produces a more realistic null expectation and avoids movement through inaccessible areas that could bias estimates of random co-occurrence (Hanks et al., 2017). The realism of null trajectories could be further improved using movement models that represent longer-term persistence, behavioural states or other ecological constraints (McClintock & Michelot, 2018; Nathan et al., 2008). Although we used a first-order vector autoregressive model for computational efficiency, the framework can accommodate alternative movement models. It is also scalable to large datasets: more than 1,500 associations were evaluated automatically without visual inspection. Because inference is conducted for individual association events rather than complete trajectories, the approach is transferable to other taxa for which concurrent animal and vessel movement data are available. With adaptations to identify potential exposure events before close proximity occurs, the simulation framework could also be extended to assess responses such as vessel avoidance (Burger et al., 2019; Fliessbach et al., 2019).

The sensitivity analysis showed that the simulation-based framework was substantially less sensitive to variation in spatiotemporal parameters than conventional threshold-defined associations. Association rates increased markedly with broader proximity thresholds and were moderately affected by minimum event duration, whereas the classification of attraction and, particularly, following remained comparatively stable across the parameter space examined. This suggests that, once a candidate association is identified, behavioural inference is driven primarily by the observed movement response rather than the precise definition of the initial proximity event, reducing dependence on arbitrary parameter choices. Similarly, varying the temporal offset used to initialize simulated trajectories had little effect on the probability of detecting attraction, indicating robustness to moderate changes in the initialisation period. Nevertheless, thresholds should be tailored to the study system and informed, where possible, by species-specific behavioural observations, experimental evidence, sensitivity analyses or calibration against independent data.

Previous studies have used auxiliary behavioural information, such as time spent sitting on the water derived from immersion sensors, to infer vessel attendance (Cianchetti-Benedetti et al., 2018). In contrast, our framework does not require prior assignment of behavioural states using methods such as hidden Markov models or expectation-maximisation binary clustering, which often require species-specific parameterisation and may produce ambiguous classifications (Bennison et al., 2018; Florko et al., 2023). Nevertheless, species behaviour can influence candidate associations. For instance, birds spending more time stationary on the water were less likely to be classified as attracted to or following vessels, indicating that many proximity events involving stationary birds were incidental.

This pattern was particularly evident for nonfishing vessels, which dominated traffic and frequently operated in coastal waters where shearwaters rest. These vessels generated numerous proximity-based associations but relatively few directed interactions. Conversely, although fishing vessels represented only approximately 3% of vessels recorded by AIS, they accounted for 69.1% of attraction events and 74.1% of following events. Together with the greater probability of attraction and following during daylight, these patterns are consistent with the known ecology of Scopoli’s shearwaters and support the biological relevance of the simulation-based classification (Navarro-Herrero et al., 2024; Sánchez-Román et al., 2019; Soriano-Redondo et al., 2016). Local vessel density had no statistically supported effect, indicating that the overall concentration of traffic did not increase the probability of a directed response. Because this metric encompassed all vessel types and was dominated by nonfishing traffic, it may not adequately represent the availability of feeding opportunities associated specifically with aggregations of fishing vessels (Navarro-Herrero et al., 2025).

The ecological importance of fishing aggregations was also reflected in the analysis of overlapping associations. The lower prevalence of overlap among attraction and following events, particularly when both responses were detected, suggests that the strongest evidence of directed movement was less likely to arise under conditions in which several vessels represented plausible targets. This supports the discriminatory value of the framework in areas of dense vessel traffic, although it does not establish which vessel elicited a response when multiple associations remain. The predominance of fishing vessels within these overlaps may reflect the spatial aggregation of fishing activity and the tendency of seabirds to exploit groups of vessels or shared foraging opportunities rather than responding independently to a single vessel (Collet et al., 2017; Rutter et al., 2024; Soriano-Redondo et al., 2016). Under these conditions, future developments could explicitly model fishing aggregations as the relevant interaction unit rather than treating each vessel independently.

By using high-resolution Automatic Identification System (AIS) data, we characterised vessel activity across fleets and jurisdictions, extending previous studies based primarily on restricted Vessel Monitoring System (VMS) data limited to national waters (Bartumeus et al., 2010; Cianchetti-Benedetti et al., 2018; Genovart et al., 2018; Reyes-González et al., 2021; Soriano-Redondo et al., 2016). However, AIS adoption remains uneven across fleets. Although terrestrial reception and coverage of industrial fishing vessels were sufficient in our study region (March et al., 2021; Welch et al., 2022), small-scale fishery vessels <15 m are not required to carry AIS transponders and are therefore underrepresented. This limits our ability to quantify seabird-fishery interactions fully and may lead to underestimation of bycatch risk and other fisheries impacts (Cortés et al., 2017; Pott & Wiedenfeld, 2017). Small recreational vessels are similarly underrepresented because those <24 m are generally exempt from AIS requirements. Indeed, recent satellite-based surveys in an area overlapping our study region found that only approximately 8.5% of detected recreational vessels carried AIS transponders (Menéndez-Blázquez et al., 2025). AIS data may therefore also underestimate seabird exposure and response to recreational vessels. Expanding electronic monitoring across small-scale fisheries and recreational fleets would help reveal these currently overlooked interactions.

Although our framework identifies directed animal-vessel interactions, movement trajectories alone cannot reveal the specific behaviours performed during these events, such as feeding on discards or approaching baited hooks. Complementary bio-logging devices, including accelerometers (Cianchetti-Benedetti et al., 2018), light and wet-dry sensors (Carneiro et al., 2022; Darby et al., 2023; Rutter et al., 2025), bird-borne cameras (Michel et al., 2021), and radar detectors (Navarro-Herrero et al., 2024), could help characterise these fine-scale behaviours and enhance our understanding of broader animal-vessel interactions, including bycatch.

Our simulation-based framework provides a scalable way to determine whether individual animal-vessel associations reflect attraction or following rather than incidental co-occurrence. Its robustness to parameter selection and implementation in the R package *intersimR,* facilitates reproducible application across taxa, vessel activities and high-traffic marine systems. Combined with complementary biologging data, the framework can help identify the animal behaviour and vessel activities most relevant to conservation and management.

## Acknowledgements

DM and LNH acknowledge support from the CIDEGENT program of the Generalitat Valenciana (CIDEGENT/2021/058) and JGS acknowledges support from ICREA Academia. LNH was supported by Pleamar (2017/2349), and BES-2017-079874 (CGL 2016-78530-R). This research was funded by the project “Minimising bycatch of seabirds and sea turtles in West African industrial fisheries” MAVA Foundation (MAVA20210/20113/20033), by Ministerio de Ciencia, Innovación y Universidades of the Spanish Government (PID2020-117155GB-I00/AEI/10.13039/501100011033 and PID2023-146782OB-100/MICIU/AEI /10.13039/501100011033 and FEDER UE). We acknowledge support from the University of Exeter’s Advanced Research Computing facilities at Penryn in carrying out this work. We thank the Govern de les Illes Balears for permits and Andrea Soriano, Laura Zango, Fernanda Pereira, Montserrat Vanerio and many undergrad and master students for their help with the fieldwork.

## Author contribution

DM: Conceptualisation, Formal analysis, Software (R code development), Writing - original draft. LNH: Fieldwork, Validation, Writing - review & editing. JGS: Conceptualisation, Funding acquisition, Writing - review & editing.

## Data availability

Raw AIS data are available from the Balearic Islands Coastal Observing and Forecasting System (SOCIB) and EXMILE SOLUTIONS LTD (MarineTraffic) and provided to the Cavanilles Institute of Biodiversity and Evolutionary Biology (ICBiBE, University of Valencia) in the context of a data sharing agreement with SOCIB, accessed 2024-06-26. The seabird GPS tracking dataset is hosted at the Seabird Tracking Database of BirdLife International (http://seabirdtracking.org) with the following IDs: 870, 871, 1618, 1619, 1622, 1623, 1624.

## Code availability

All analyses and plots were undertaken using the R programming language (R Core Team 2019). The simulation framework is implemented in the R package *intersimR*, available at Github (https://github.com/spatialmarine/intersimR/). The following GitHub repository contains the R code to conduct the analysis presented in this work (https://github.com/dmarch/seabird-ships).

## Statement on inclusion

Our study on seabird-fishing interactions in the NW Mediterranean Sea was conducted by local scientists, ensuring alignment with regional conservation priorities. This approach, leveraging bird-borne sensors, not only enhances local seabird conservation efforts but also offers a valuable method for studying similar interactions in other regions with complex maritime traffic and/or limited vessel tracking data.

## Generative AI statement

GitHub Copilot was used during the development of the *intersimR* package to assist with code refactoring, and the conversion of existing analytical scripts into a structured R package. ChatGPT was used to improve the clarity and grammar of the manuscript text. Generative AI tools were not used to interpret scientific results, draw conclusions, or make scientific decisions. All AI-generated suggestions were reviewed, validated, and edited by the authors as appropriate. The authors take full responsibility for the content of the manuscript.

