## Supplementary Information for "A simulation-based framework to detect marine animal-vessel interactions using tracking data"

### Contents

Supplementary Methods (Supplementary Methods 1-2)

Supplementary Tables (Supplementary Tables 1-5)

Supplementary Figures (Supplementary Figures 1-4)

Supplementary References

### Supplementary Methods

**Supplementary Methods S1. Selection of the pre-association offset for attraction tests**

To evaluate attraction behaviour, simulations were initialised at a fixed temporal offset prior to the onset of each association event (see Section 2.3.2 of the main manuscript). The duration of this pre-association window must balance two constraints: (i) it should be sufficiently long to capture potential directed movement towards vessels, and (ii) it should remain behaviourally connected to the focal association event.

For Scopoli's shearwaters, we selected a 30-min offset prior to the observed onset of association. This choice is consistent with the movement capabilities of the species and expected sensory detection ranges of vessels. Assuming typical flight speeds of approximately 30-50 km $h^{-1}$, individuals can travel 15-25 km within 30 minutes, which falls within reported detection distances of fishing vessels (≈10-30 km) (Skov and Durinck 2001; Orben et al. 2021). This supports the assumption that behavioural responses leading to attraction are likely to occur within this temporal window.

To further assess whether a 30-min offset reflects realistic response times, we quantified the elapsed time between the initial encounter with a vessel and the subsequent onset of association. The resulting distribution was strongly right-skewed, with most values occurring within a few hours (Figure S1a), indicating that response times on the order of tens of minutes are plausible. Similarly, elapsed times between consecutive association events showed a comparable distribution (Figure S1b), suggesting that longer offsets would increase the likelihood of initiating simulations while the animal was still engaged in a previous interaction. This could confound the evaluation of attraction because the simulated trajectory might be initialized while the bird was still responding to a previous vessel association.

Because the choice of offset may influence the behaviour of the null model, we evaluated its sensitivity by repeating the attraction test under alternative offset durations (see Supplementary Methods S2b). This analysis confirmed that the main conclusions were robust to reasonable variation in the offset parameter.

**Supplementary Methods S2. Sensitivity analysis of the simulation approach**

We performed two complementary sensitivity analyses to evaluate the robustness of the simulation-based inference approach and to assess how parameter choices influenced the detection of interaction events.

**(a) Sensitivity to spatio-temporal parameters**

To evaluate the sensitivity of detected interaction events to spatio-temporal parameter selection, we explored a factorial combination of three proximity distance thresholds (1, 1.5, and 3 km), three minimum event durations (10, 15, and 20 min), and three maximum temporal gaps allowed between consecutive events (10, 30, and 60 min), resulting in 27 parameter combinations. Due to the computational demands associated with repeatedly running the simulation framework across all parameter combinations, analyses were conducted using a random subset of 50 foraging trips selected from the full dataset of 2,686 trips. Trips were randomly sampled using a fixed random seed to ensure reproducibility. The temporal offset used to initialise simulated trajectories for the attraction test was kept constant at 30 min before the start of the association event.

For each trip and parameter combination, we calculated the number of detected interaction events for each interaction type: associations identified using the threshold-based approach, and attraction and following events identified as significant by the simulation framework. Interaction rates were analysed using a generalized linear mixed model (GLMM) fitted with a negative binomial distribution and log link using the *glmmTMB* package. The response variable corresponded to the number of detected events per trip and interaction type. Fixed effects included interaction type, proximity distance threshold, minimum duration threshold, and maximum temporal gap threshold, as well as their interactions with interaction type. Foraging trip identity was included as a random intercept to account for repeated measurements across parameter combinations, and the log-transformed trip duration (days) was incorporated as an offset term to standardize event rates by foraging effort.

Model diagnostics were evaluated using simulated residuals generated with the *DHARMa* package. Residual diagnostics indicated no evidence of residual overdispersion (dispersion test: $p = 0.092$) or zero inflation (zero-inflation test: $p = 0.808$). Although the Kolmogorov-Smirnov test detected a significant deviation from residual uniformity ($p < 0.001$), visual inspection of residual distributions and residual patterns across predictor levels revealed only minor and localized deviations from uniformity, primarily associated with extreme parameter values. Overall, the model was considered to provide an adequate representation of the data.

#### (b) Sensitivity to time offset duration in the attraction test

We conducted an additional sensitivity analysis to evaluate the influence of the temporal offset used to initialise simulated trajectories in the attraction test. The offset duration, corresponding to the period preceding the start of the association used to initialise simulations, was varied between 10 and 60 min in 10-min increments, while keeping all spatio-temporal thresholds constant (proximity distance = 1.5 km, minimum duration = 15 min, maximum temporal gap = 30 min).

Due to the computational demands associated with repeatedly running simulations across offset values, analyses were conducted using a subset of 50 association events randomly selected from the full dataset of association events (n = 1,567). To ensure balanced representation of both outcomes, the subset was stratified according to the original attraction test result, including 25 significant and 25 non-significant associations under the reference parameter configuration. Each association event was subsequently reanalysed using the different offset values. For each event and offset combination, the attraction test produced a binary outcome (1 = significant attraction event, 0 = non-significant event), which was analysed using a binomial generalized linear mixed model (GLMM) with offset duration included as a continuous fixed effect and association identity included as a random intercept.

### Supplementary Tables

**Supplementary Table S1.** Description of explanatory variables included in the GLMMs testing factors influencing the probability that candidate associations were classified as attraction or following.

| ***Variable*** | ***Type*** | ***Description*** | ***Estimation / Data source*** |
|---|---|---|---|
| Vessel type | Categorical | Type of vessel involved in the association (i.e. fishing, nonfishing) | Classified based on vessel information from AIS messages and two different fishing vessel registers, the EU Fishing Fleet Register and the Global Fishing Watch. |
| Seabird stationary behaviour | Continuous (0–100%) | Percentage of time the seabird remains stationary during the association event | Computed as the proportion of GPS fixes classified as "resting" within the duration of the event, based on EMbC algorithm (Garriga et al. 2016). |
| Day/night phase | Categorical (day / night) | Light condition during the association event. | Determined from the time and geographic position of the event using the *suntools* package in R. |
| Vessel density (30 km radius) | Continuous | Number of vessels detected within a 30 km radius around the association during the event. | Estimated from AIS data by counting unique vessel MMSI within a 30 km buffer and concurrent time window. |

**Supplementary Table S2.** Summary of overlapping associations across association classes. Overlapping associations were defined as events in which the same individual associated with more than one vessel during overlapping temporal windows. Fishing-only overlaps refer to overlapping events where both the focal and overlapping vessels were classified as fishing vessels. The number of overlapping vessels refers to additional vessels overlapping with the focal event.

| *Association type* | *Association events* | Events overlapping another association | *Fishing-only overlaps* | *Overlapping vessels* |
|---|---|---|---|---|
| | *N* | *N (%)* | *%* | *mean (range)* |
| Association | 1,567 | 441 (28.1%) | 66.7 | 1.7 (1–9) |
| Attraction | 698 | 169 (24.2%) | 67.5 | 1.6 (1–7) |
| Following | 432 | 82 (18.9%) | 65.9 | 1.6 (1–9) |
| Attraction and Following | 291 | 39 (13.4%) | 61.5 | 1.4 (1–7) |

**Supplementary Table S3.** Results of the generalized linear mixed model (GLMM) with a negative binomial distribution assessing the effects of spatio-temporal parameter thresholds on interaction rates across interaction types (association, attraction, and following). Incidence rate ratios (IRR) and 95% confidence intervals (CI) are provided for each predictor, indicating the multiplicative change in predicted interaction rates relative to the reference categories. Reference levels correspond to association events, proximity distance = 1 km, minimum duration = 10 min, and maximum temporal gap = 10 min. Interaction terms indicate how the effects of each parameter differed for attraction and following events relative to association events. Foraging trip identity (trip ID) was included as a random intercept to account for repeated measurements across parameter combinations.

| *Predictors* | *Incidence Rate Ratios* | *CI* | *p* |
|---|---|---|---|
| (Intercept) | 0.02 | 0.01 – 0.05 | **<0.001** |
| $\text{Method}_{\text{Attract}}$ | 0.47 | 0.27 – 0.83 | **0.009** |
| $\text{Method}_{\text{Follow}}$ | 0.26 | 0.12 – 0.53 | **<0.001** |
| $\text{Proximity}_{1.5}$ | 2.22 | 1.57 – 3.14 | **<0.001** |
| $\text{Proximity}_{3}$ | 17.88 | 13.11 – 24.39 | **<0.001** |
| $\text{Duration}_{15}$ | 0.59 | 0.50 – 0.69 | **<0.001** |
| $\text{Duration}_{20}$ | 0.36 | 0.30 – 0.44 | **<0.001** |
| $\text{Time gap}_{30}$ | 1.02 | 0.86 – 1.22 | 0.814 |
| $\text{Time gap}_{60}$ | 1.06 | 0.89 – 1.26 | 0.548 |
| $\text{Method}_{\text{Attract}} \times \text{Proximity}_{1.5}$ | 0.50 | 0.27 – 0.92 | **0.026** |
| $\text{Method}_{\text{Follow}} \times \text{Proximity}_{1.5}$ | 0.52 | 0.26 – 1.02 | 0.058 |
| $\text{Method}_{\text{Attract}} \times \text{Proximity}_{3}$ | 0.22 | 0.13 – 0.37 | **<0.001** |
| $\text{Method}_{\text{Follow}} \times \text{Proximity}_{3}$ | 0.07 | 0.04 – 0.14 | **<0.001** |
| $\text{Method}_{\text{Attract}} \times \text{Duration}_{15}$ | 1.45 | 0.98 – 2.15 | 0.061 |
| $\text{Method}_{\text{Follow}} \times \text{Duration}_{15}$ | 1.70 | 0.94 – 3.08 | 0.078 |
| $\text{Method}_{\text{Attract}} \times \text{Duration}_{20}$ | 1.83 | 1.19 – 2.79 | **0.006** |
| $\text{Method}_{\text{Follow}} \times \text{Duration}_{20}$ | 2.75 | 1.51 – 5.03 | **0.001** |

| | | | |
|---|---|---|---|
| $Method_{Attract}$ × Time $gap_{30}$ | 0.96 | 0.64 – 1.46 | 0.860 |
| $Method_{Follow}$ × Time $gap_{30}$ | 0.84 | 0.44 – 1.62 | 0.607 |
| $Method_{Attract}$ × Time $gap_{60}$ | 1.02 | 0.68 – 1.53 | 0.940 |
| $Method_{Follow}$ × Time $gap_{60}$ | 1.49 | 0.84 – 2.66 | 0.172 |
| **Random Effects** | | | |
| $\tau_{00\ tripID}$ | 5.65 | | |
| ICC | 0.86 | | |
| Marginal $R^2$ / Conditional $R^2$ | 0.166 / 0.885 | | |

**Supplementary Table S4.** Summary of fixed and random effects from the generalized linear mixed model (GLMM) assessing the effect of the temporal offset used to initialise simulated trajectories on the probability of detecting a significant attraction event. Odds ratios (OR) and 95% confidence intervals (CI) are provided for the fixed effect, with values indicating the relative change in the odds of classifying an association as a significant attraction event. Random effects represent variation among association events (association ID).

| *Predictors* | *Odds Ratios* | *CI* | *p* |
|---|---|---|---|
| (Intercept) | 1.26 | 0.24 – 6.71 | 0.790 |
| Time offset | 1.01 | 0.99 – 1.03 | 0.502 |
| **Random Effects** | | | |
| $\tau_{00\ \mathrm{associationID}}$ | 26.43 | | |
| ICC | 0.89 | | |
| Marginal $R^2$ / Conditional $R^2$ | 0.000 / 0.889 | | |

**Supplementary Table S5.** Summary of fixed and random effects from two independent generalized linear mixed models (GLMMs) .assessing factors influencing the probability that an association was classified as significant attraction or following. Odds ratios (OR) and 95% confidence intervals (CI) are provided for fixed effects, with values indicating the relative likelihood of a significant interaction for each predictor. Random effects represent variation among individuals (organism ID). See Supplementary Table S1 for variable descriptions.

| | Attraction test | | | Following test | | |
|---|---|---|---|---|---|---|
| ***Predictors*** | ***Odds Ratios*** | ***CI*** | ***p*** | ***Odds Ratios*** | ***CI*** | ***p*** |
| (Intercept) | 1.99 | 1.66 – 2.39 | **<0.001** | 0.86 | 0.70 – 1.07 | 0.170 |
| Vessel type [Nonfishing] | 0.83 | 0.65 – 1.06 | 0.129 | 0.59 | 0.45 – 0.78 | **<0.001** |
| Stationary behaviour | 0.97 | 0.97 – 0.98 | **<0.001** | 0.97 | 0.97 – 0.98 | **<0.001** |
| Time of day [night] | 0.66 | 0.51 – 0.84 | **0.001** | 0.68 | 0.50 – 0.91 | **0.011** |
| Vessel density | 0.94 | 0.84 – 1.05 | 0.270 | 0.90 | 0.79 – 1.03 | 0.111 |
| Random Effects | | | | | | |
| $\tau_{00\ organismID}$ | 0.06 | | | 0.24 | | |
| ICC | 0.02 | | | 0.07 | | |
| Marginal $R^2$ / Conditional $R^2$ | 0.234 / 0.249 | | | 0.226 / 0.278 | | |

### Supplementary Figures

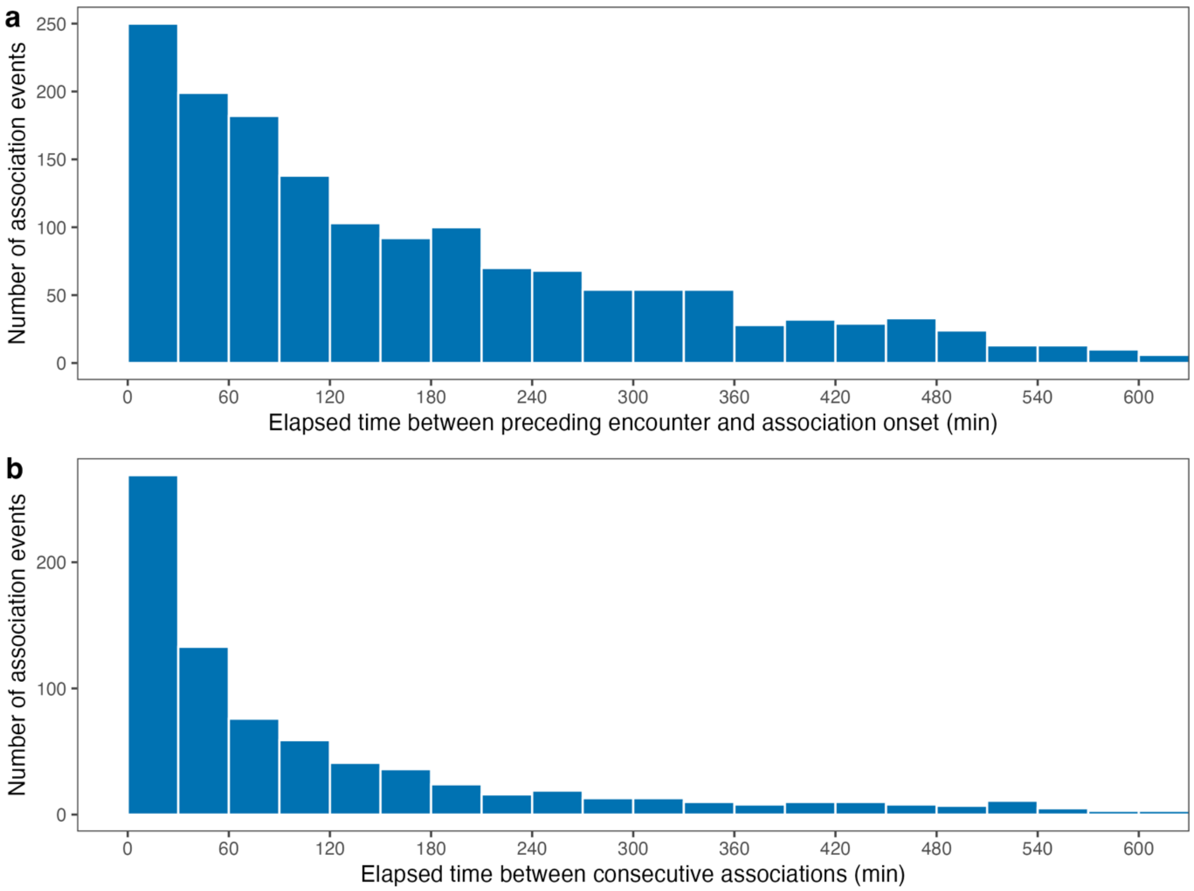


**Supplementary Figure S1.** Distribution of elapsed times between seabird-vessel interaction events. (**a**) Distribution of elapsed times between the onset of each association event and the start of the preceding encounter with the same vessel, shown in 30-min intervals (n = 1,567). (**b**) Distribution of elapsed times between consecutive seabird-vessel associations across all birds and trips, shown in 30-min intervals. Includes only associations with a preceding event within the same foraging trip (n = 1,055). For clarity, values >600 min are not shown in both plots.

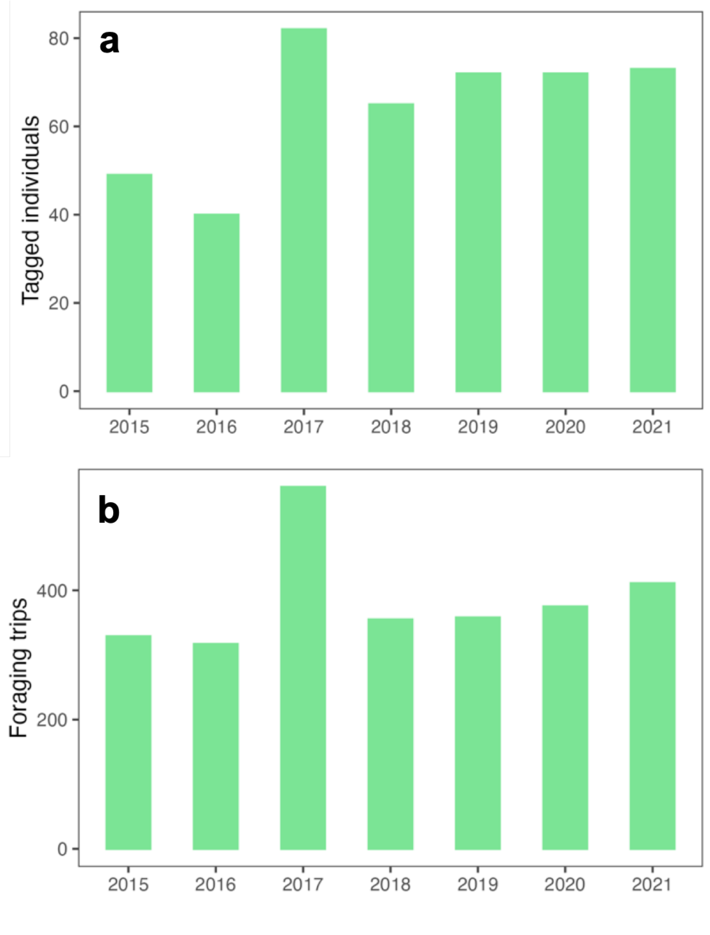


**Supplementary Figure S2.** Summary of the seabird tracking data. (**a**) Number of tagged individuals per year. (**b**) Number of foraging trips per year.

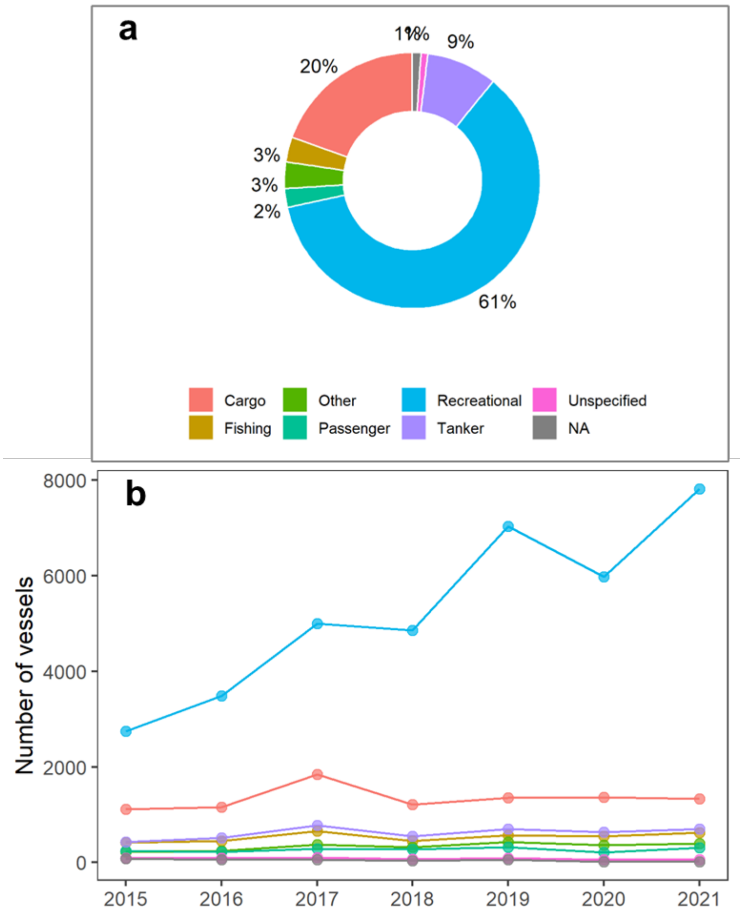


**Supplementary Figure S3.** Overview of the vessel tracking data. (**a**) Proportional contribution of each vessel category across all years combined (n = 28,253 unique vessels). (**b**) Annual number of vessels recorded in each category between 2015 and 2021.

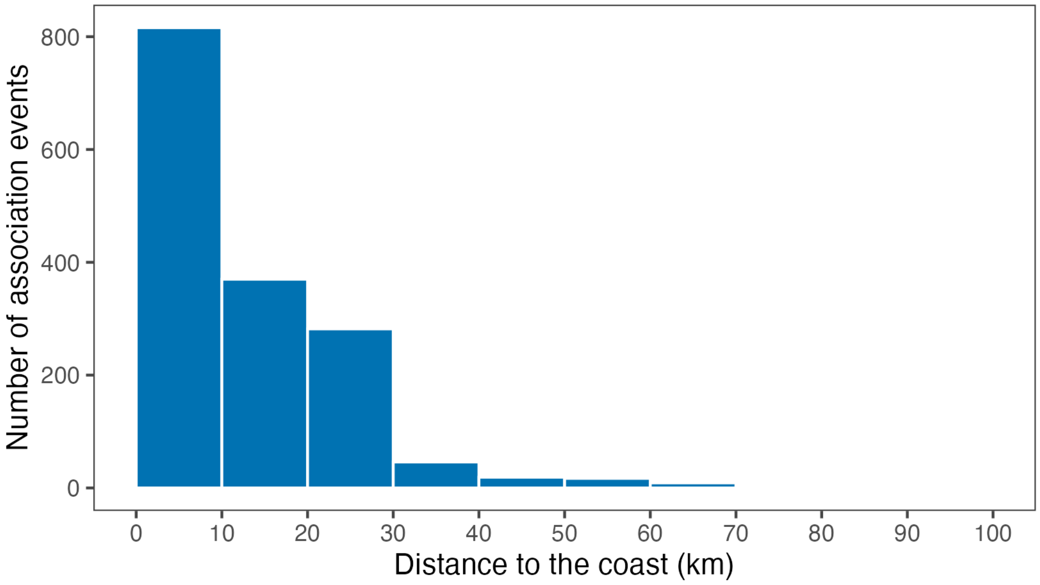


**Supplementary Figure S4.** Distribution of association events as a function of distance to the coast, shown in 10-km intervals. For clarity, distances >100 km are omitted.
